# Sex- and Cell Type-Specific Mitochondrial Drug Responses in Patient-Derived Models of Sporadic Alzheimer’s Disease

**DOI:** 10.64898/2026.11.19.767925

**Authors:** Daniel Chen, Maggie Benson, Vivien Csikos, Ashley Tetlow, Vaidehi Bhavaraju, Keith Smith, Heather M. Wilkins

## Abstract

Alzheimer’s disease (AD) is associated with mitochondrial dysfunction, yet therapeutic discovery has often overlooked cell-type-specific vulnerabilities and biological sex. We performed a high-throughput screen of a mitochondrial-targeted library in induced pluripotent stem cell (iPSC)-derived neurons and astrocytes from AD and age/sex-matched non-demented (ND) donors. Oxygen consumption rate (OCR) was used as the primary functional readout, with screening performed separately in male and female cells. Male and female AD neurons exhibited impaired mitochondrial respiration across complexes I–IV, whereas astrocytes displayed divergent phenotypes characterized by reduced respiration in males and increased respiration in females. Drug screening results varied by cell type and sex. P38 MAPKI restored mitochondrial respiration in both male and female AD neurons and was associated with reduced mitochondrial oxidative stress and normalization of mitochondrial mass in male neurons. P38 MAPKI additionally reduced secreted Aβ_42_ and phospho-tau (Thr217). In astrocytes, anthocyanidin selectively restored respiration in female AD cells without substantial changes in mitochondrial mass, membrane potential, or oxidative stress, whereas PPARα agonist selectively rescued mitochondrial mass in male AD astrocytes and was associated with increased antioxidant signaling. Proteomic profiling demonstrated distinct molecular states across neurons and astrocytes and revealed that functional mitochondrial rescue did not necessarily require normalization of mitochondrial pathway signatures. These findings demonstrate that mitochondrial dysfunction and therapeutic response in AD are shaped by both cell type and biological sex. Incorporating these variables into drug discovery may enable more precise identification of therapeutic strategies tailored to distinct cellular states in AD.

## Introduction

Alzheimer’s disease (AD) is a progressive neurodegenerative disorder characterized by accumulation of amyloid-β (Aβ) plaques and hyperphosphorylated tau (p-tau) tangles, with progressive memory impairment and cognitive decline [1]. Approximately 5% of all AD cases are associated with known pathogenic variants in genes including *Amyloid Precursor Protein (APP)* and *Presenilin ½ (PSEN1/2)* [2]. However, many cases lack a clearly defined etiology and are considered sporadic. Sporadic AD cases are thought to arise from complex interactions between aging, genetic susceptibility, metabolism, inflammation, and other environmental factors [3]. Because AD is multifactorial and heterogeneous, it presents a major challenge for therapeutic development.

Despite decades of research, treatment options for AD remain limited. Classical treatment options including Donepezil, Rivastigmine, and Memantine primarily provide symptomatic relief without substantially altering disease progression [4]. More recently monoclonal antibodies (mAbs) targeting amyloid plaques have gained clinical approval; however, meta-analyses and clinical observations suggest that these therapies provide only modest cognitive benefit [5, 6]. Moreover, mAbs carry the risk of severe adverse side effects, such as amyloid-related imaging abnormalities (ARIA), which can manifest as cerebral edema and hemorrhage [6, 7]. Clinical studies involving mAbs restrict patients with microhemorrhages, superficial siderosis, or cerebral amyloid angiopathies; therefore, the real-world risk of these drugs may differ [8, 9]. Together, these findings underscore the need for alternative therapeutic strategies that target disease mechanisms beyond amyloid deposition alone.

The emergence of induced pluripotent stem cell (iPSC) technology has enabled the development of patient-derived cellular systems that can model AD biology within a human context [10]. iPSC-derived brain cells retain patient-specific genetic background and exhibit disease relevant phenotypes including amyloid and tau production, mitochondrial dysfunction, altered metabolism, oxidative stress, and inflammatory signaling [11, 12, 13, 14]. Importantly, these models provide an opportunity to investigate complex cellular interactions and multifactorial mechanisms that would be difficult to recapitulate in conventional experimental models.

To date, many iPSC-based studies have focused primarily on neurons [10]. But increasing evidence suggests that non-neuronal cell types, including astrocytes, play an essential role in disease onset and progression. Astrocytes are critical regulators of neuronal homeostasis and function [15]. Dysfunctional astrocytes can exacerbate neuronal stress through cell-autonomous and non-cell-autonomous mechanisms contributing to neurodegeneration and disease propagation [15, 16]. Given the high metabolic interdependence between astrocytes and neurons, therapeutic targets identified in neuron-only systems may overlook the important contributions from glial cells and their intercellular metabolic coupling [17]. Therefore, incorporating both neuronal and astrocyte models may provide a more physiologically relevant framework for drug discovery in AD.

Among the many pathological mechanisms that are implicated in AD, mitochondrial dysfunction has emerged as a promising therapeutic avenue [18]. Mitochondria play a central role in cellular bioenergetics, oxidative stress response, and inflammatory signaling [18, 19]. Defects in mitochondrial respiration, dynamics, and reactive oxygen species (ROS) production and handling, have all been observed early in AD progression [18, 20]. Clinical imaging studies using 2-deoxy-2-[fluorine-18]fluoro-D-glucose position emission tomography (FDG-PET) and functional magnetic resonance imaging (fMRIs) further demonstrate that metabolic abnormalities may precede overt cognitive decline [21]. Furthermore, mitochondrial dysfunction interacts bidirectionally with both amyloid and tau pathology [18, 22], forming a vicious cycle that promotes neuronal injury and disease progression. These findings suggest that mitochondrial dysfunction may represent an early and targetable aspect of AD.

An additional challenge in AD research is the underrepresentation of biological sex, despite well-established sex differences in disease prevalence, progression, and therapeutic response [23, 24]. Women account for nearly two-thirds of all AD cases and exhibit distinct molecular and clinical features compared to men [25]. Emerging evidence further suggests that mitochondrial function, oxidative stress responses, and neuroinflammatory signaling may differ between males and females in AD [26]. Incorporating sex as a biological variable may improve the identification of more effective and personalized treatment strategies.

In this study, we leveraged a high-throughput drug screen approach using AD iPSC-derived neurons and astrocytes to identify compounds capable of restoring mitochondrial function. Our approach aimed to capture disease-relevant mechanisms that are often overlooked in traditional AD models by integrating patient-derived multicellular models with mitochondrial phenotypic screening, while factoring in sex as a biological variable. Through this study, we sought to identify novel or repurposed therapeutic candidates capable of targeting shared metabolic vulnerabilities across multiple brain cell types in AD.

## Results

### High-throughput mitochondrial drug screen identified four protective hits in female iPSC-derived neurons

To identify compounds capable of restoring mitochondrial function in AD, we performed a high-throughput drug screen of compounds that target mitochondrial function using iPSC-derived neurons and astrocytes generated from sporadic AD patients and non-demented donors (ND). Our mitochondrial-targeted library was composed of 583 compounds with various mechanisms of action. Oxygen consumption rate (OCR), measured via Seahorse analysis, served as a primary readout, with protective hits defined as compounds that restored AD cell OCR to within ±20% of the mean ND control, based on fold change. The list of drugs and classifications are shown in **Table 1**, **Figure 1**.

**Figure 1.**
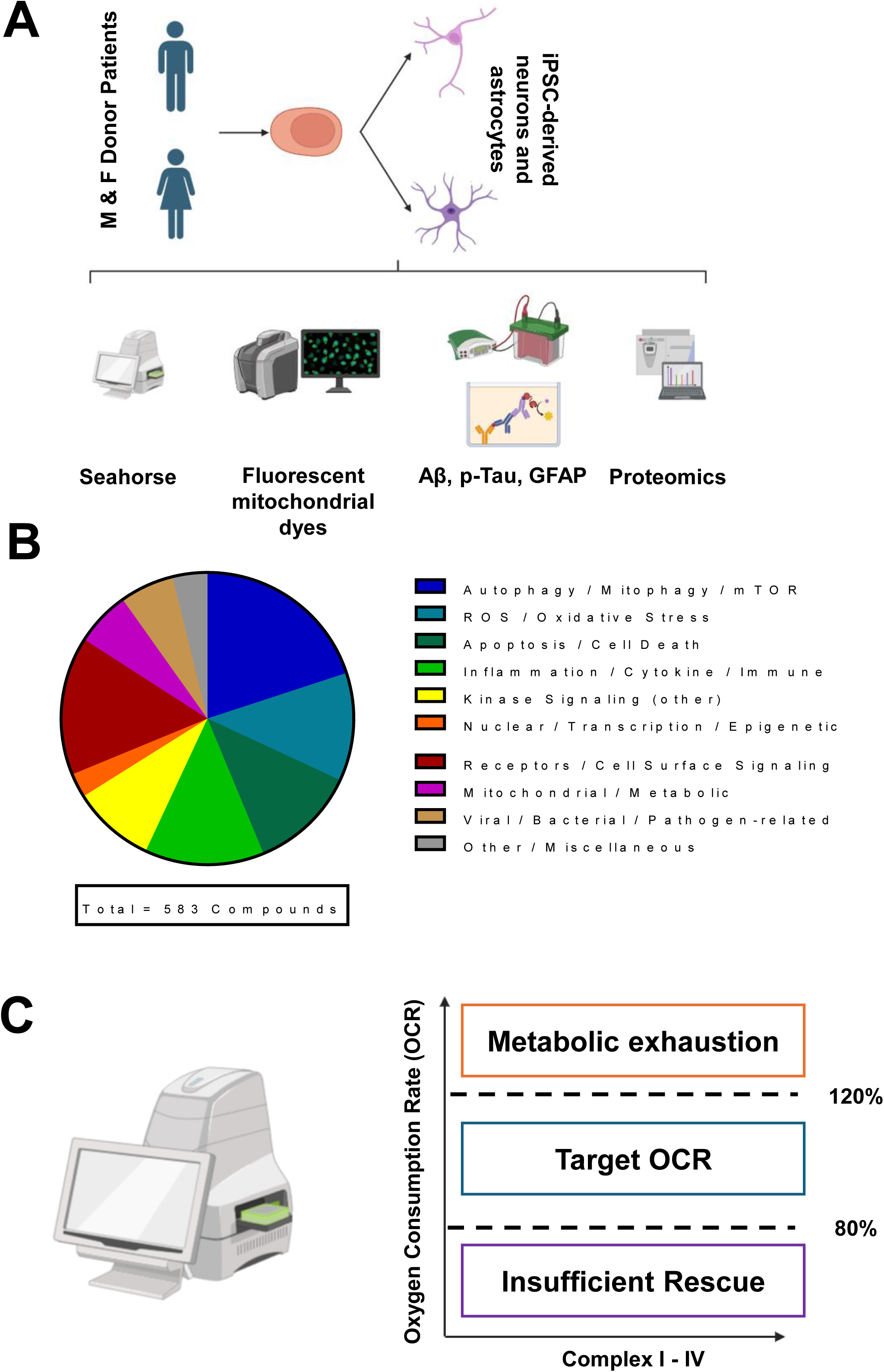

### Mitochondrial drug screen identified one protective hit in iPSC-derived neurons

We demonstrate sex-specific differences in OCR in iPSC-derived neurons across complexes I-IV, measured using the seahorse electron transport chain (ETC) flux assay (**Figure 2A**). Both male and female neurons showed significant reductions in complex I-IV OCR (multiple unpaired t-test with Welch correction Holm-Sidak method, adjusted p value <0.000001; **Figure 2A**). Screening of female AD neurons identified four compounds that met the ±20% OCR restoration criterion: PPARγ agonist, diamine oxidase, P38 MAPKI, and neuronal nitric oxide synthase (**Figure 2B-E**). In male AD neurons, three compounds met the primary hit threshold, PPARγ agonist, P38 MAPKI, and an essential amino acid. PPARγ agonist and P38 MAPKI were identified as hits in both sexes, suggesting that these compounds target mitochondrial pathways impaired across the AD patient population regardless of sex (**Figure 2F-I**). In contrast, an essential amino acid was a male-specific hit, while diamine oxidase and neuronal nitric oxide synthase were female specific hits, indicating sex-specific mechanisms of mitochondrial vulnerability in sporadic AD neurons.

**Figure 2.**
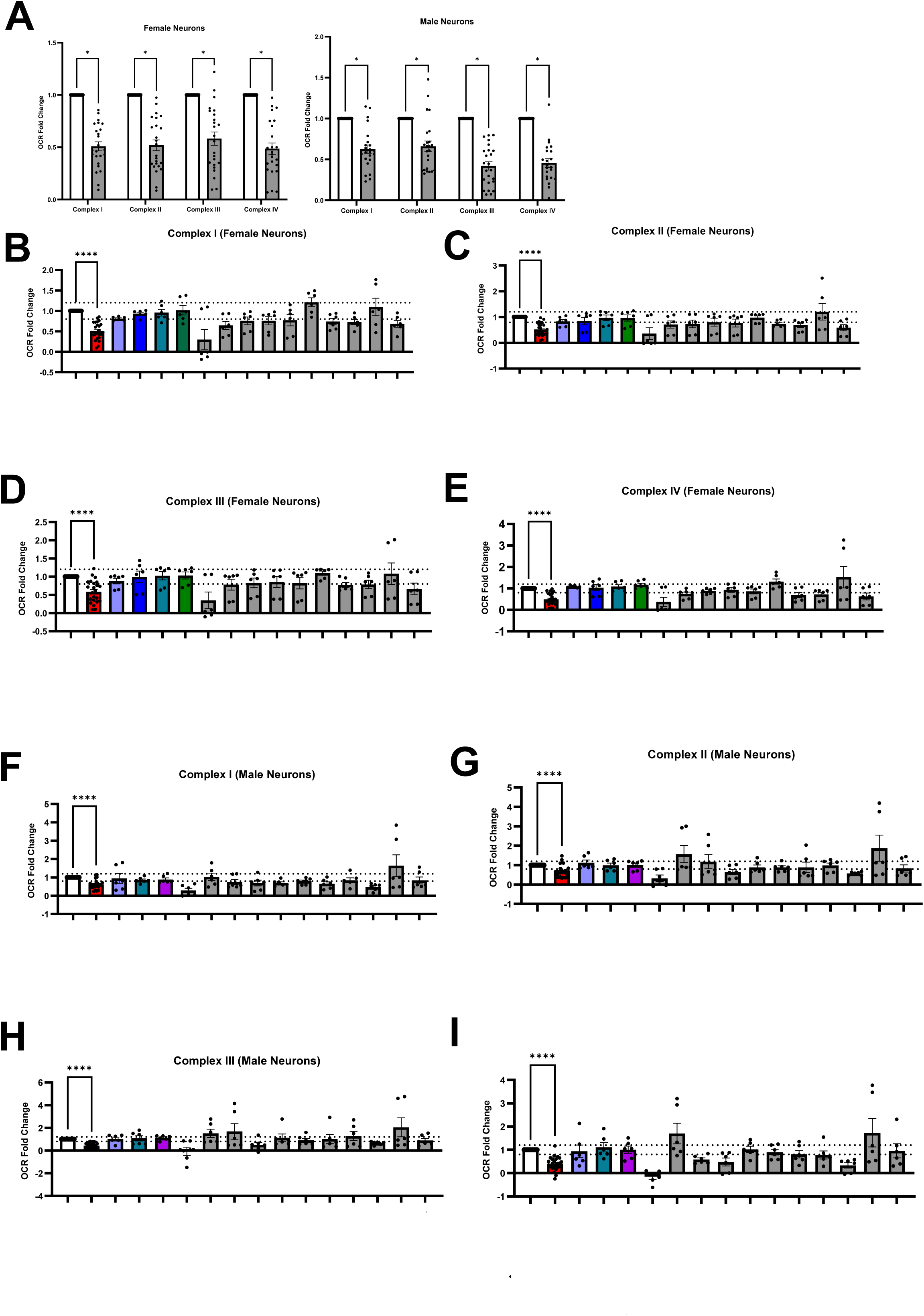

In female AD iPSC-derived neurons MitoTracker fluorescence intensity did not differ substantially across groups (**data not shown**). However, MitoSOX fluorescence was significantly elevated in female AD neurons compared to ND controls (**Figure 3A**; Ordinary one-way ANOVA with multiple comparisons to AD, α = 0.05; adjusted p-value = 0.0279). Treatment with the drug hits for female neurons; PPARγ agonist, diamine oxidase, P38 MAPKI, and neuronal nitric oxide synthase, individually reduced MitoSOX signal toward ND levels, with only P38 MAPKI showing statistical significance (**Figure 3A**; one-way ANOVA with multiple comparisons α = 0.05; adjusted p-value = 0.0499). TMRE-based membrane potential measurements showed modest and variable differences between AD and ND groups with drug-treated cells showing intermediate values (**data not shown**). This pattern is consistent with reports that MMP compensation can mask mitochondrial dysfunction in iPSC-based AD models [27].

**Figure 3.**
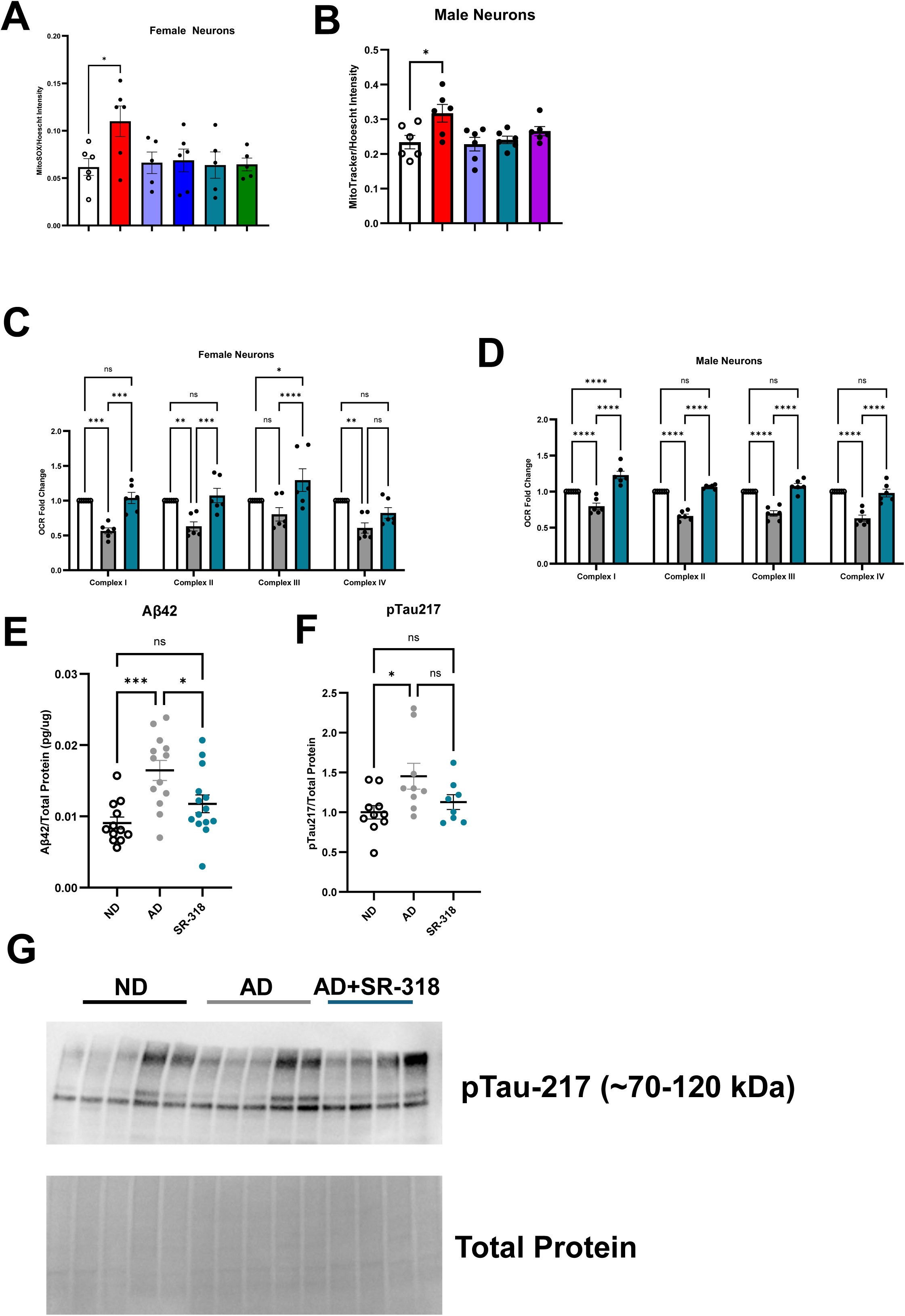

In male AD iPSC-derived neurons MitoTracker fluorescence was significantly elevated in AD neurons compared to ND controls (**Figure 3B**; Ordinary one-way ANOVA with multiple comparisons to AD, p = 0.0136), suggesting altered mitochondrial accumulation or impaired mitochondrial turnover. Treatment with PPARγ agonist and P38 MAPKI both normalized MitoTracker levels comparable to ND controls (**Figure 3B**; Ordinary one-way ANOVA with multiple comparisons to AD; P_PPARγ agonist_ = 0.0078, p_P38 MAPKI_ = 0.0245). Male AD neurons showed no change in MitoSOX or TMRE fluorescence compared to ND controls (**data not shown**). Our secondary screen confirmed P38 MAPKI normalized OCR in both male and female iPSC neurons across multiple independent donors (**Figure 3C,D** n=6).

### P38 MAPKI reduces Aβ_42_ secretion and tau hyperphosphorylation in AD iPSC-derived neurons

Given its consistent multi-parameter mitochondrial rescue profile – including OCR and ROS production, P38 MAPKI was selected as the lead neuronal hit for downstream analysis and characterization. To determine whether P38 MAPKI treatment also affected canonical AD pathological markers, we measured secreted Aβ_42_ levels (the major component of amyloid plaques; [28]) by ELISA and phosphorylated tau (pTau) abundance by western blotting for pTau-217 (widely considered to be the most specific phosphorylated tau isoform for AD; [29]). Consistent with the disease phenotype of AD, Aβ_42_ levels normalized to total protein were significantly elevated in AD iPSC-derived neurons (**Figure 3E**; Ordinary one-way ANOVA with multiple comparisons to ND, p = 0.0003) relative to ND controls in both sexes, and AD iPSC neurons were rescued with P38 MAPKI treatment. Furthermore, pTau levels normalized to total protein were significantly elevated in AD iPSC neurons compared to ND controls (**Figure 3F, G**; Ordinary one-way ANOVA with multiple comparisons to ND, p = 0.0005). P38 MAPKI treatment substantially reduced pTau-217 abundance in AD iPSC neurons.

### P38 MAPKI alters expression of mitochondrial electron transport chain components in iPSC derived neurons from sporadic AD patients

We performed global data-independent acquisition (DIA) proteomic profiling of iPSC-derived neurons and astrocytes. Neurons were evaluated with and without treatment with P38 MAPKI, whereas astrocytes were analyzed with and without their sex-specific treatment. Across all samples, DIA proteomics quantified approximately 10,000 proteins per cell-type, enabling pathway-level interrogation of disease-associated and treatment-induced molecular changes. Consistent with previous observations from functional mitochondrial assays, we observed unique proteomic remodeling between neurons and astrocytes, suggesting cell-type-specific responses in AD.

Examining mitochondrial ETC subunit abundance identified upregulation across complex I, II, II, IV and V components in female AD iPSC neurons compared to ND neurons (**Figure 4 A, C, E, G, I**). Treatment with P38 MAPKI in female AD iPSC neurons largely normalized expression of mitochondrial ETC components with some variability noted (**Figure 4 A, C, E, G, I**), for example NDUFAF5 and NDUFAF6 were upregulated further with treatment, complex III components were upregulated with treatment as well, with mixed outcomes in complex IV and V components for female AD neurons were observed.

In male AD iPSC neurons compared to ND neurons ETC components were largely downregulated (**Figure 4 B, D, F, H, J**). Treatment with P38 MAPKI in male AD neurons showed similar effects to what was observed in female AD neurons with respect to ETC component abundance, specific components were normalized while others did not change. Complex I components NDUFAF5 and NDUFAF6 were increased by P38 MAPKI treatment in male AD neurons (**Figure 4B**). Complex II components SDHD and SDHAF1; Complex III components UQCR10, UQCC1, and UQCC3; Complex IV components mtCO1, mtCO2, XOA6, COX18, COX7B; and complex V components ATP5F1B, ATP5F1C, and ATP5MC were also normalized in male AD neurons treated with P38 MAPKI (**Figure 4D,F,H,J**).

### p38 MAPK is significantly elevated in AD iPSC-derived neurons

We measured total phosphorylated p38 MAPK (phosphor-Thr180/Tyr182) normalized to total protein in ND, AD, and P38 MAPKI-treated neurons to assess the activity state of the p38 MAPK pathway in iPSC-derived neurons. Phospho-p38 levels were significantly elevated in AD neurons (**Supplemental Figure 1**; Ordinary one-way ANOVA with multiple comparisons to ND, p = 0.0197) relative to ND controls. These data indicate an increase in p38 MAPK pathway activity in sporadic AD neurons. As expected, P38 MAPKI treatment did not alter phospho-p38 abundance, which reflect its mechanism of action as an inhibitor of further downstream phosphorylation by active p38 α and β, without affecting the upstream mitogen-activated protein kinase kinase 3/6 (MKK3/6)-mediated phosphorylation of p38 itself. These findings confirm that p38 MAPK is aberrantly activated in sporadic AD iPSC neurons and that P38 MAPKI acts downstream of this activation event to inhibit substrate-level signaling.

### P38 MAPKI alters expression of mitochondrial quality control protein in iPSC derived neurons from sporadic AD patients

P38 MAPKI treatment also normalized mitochondrial dynamic pathway protein expression in both male and female iPSC neurons (**Figure 5A, B**). P38 MAPKI treatment upregulated SIRT3 expression in both male and female AD iPSC neurons and had mixed effects on other regulators of mitochondrial biogenesis (**Figure 5C, D**). Mitophagy pathway proteins were largely down regulated in female iPSC AD neurons, which was normalized by P38 MAPKI treatment (**Figure 5E**). Male neurons showed largely upregulated expression of mitophagy proteins with mixed effects from P38 MAPKI treatment (**Figure 5F**). P38 MAPKI treatment upregulated mitochondrial protease expression in female AD iPSC neurons with mixed effects in male AD iPSC neurons (**Figure 5G, H**).

### P38 MAPKI alters mitochondrial pathway expression in iPSC derived neurons from sporadic AD patients

Female AD neurons showed downregulation of mitochondrial dysfunction, nerve growth factor pathway, and upregulation of complex I biogenesis, oxidative phosphorylation, and respiratory electron transport chain (**Figure 6A**). P38 MAPKI treatment upregulated mitochondrial protein import, mitochondrial protein degradation, and mitochondrial translation pathways in female AD neurons (**Figure 6B**).

Male AD neurons showed down regulation of complex I biogenesis with upregulation of autophagy and deubiquitylation (**Figure 6C**). P38 MAPKI treatment upregulated synaptogenesis pathways and protein sorting pathways (**Figure 6D**). Other pathway changes can be found within **Supplemental Tables 1** and **2**.

### Mitochondrial drug screen identified distinct protective hits in female and male iPSC-derived astrocytes

Male AD astrocytes displayed a significant decrease in OCR (multiple unpaired t-test with Welch correction Holm-Sidak method, adjusted p-value_complex I_ = 0.000029, adjusted p-value_complex II_ < 0.000001, adjusted p-value_complex III_ = 0.000007, adjusted p-value_complex III_ = 0.000005; **Figure 7A**), but female astrocytes showed a significant increase in OCR across complex I-IV (multiple unpaired t-test with Welch correction Holm-Sidak method, adjusted p-value_complex I_ = 0.000790, adjusted p-value_complex II_ = 0.000020, adjusted p-value_complex III_ = 0.000020, adjusted p-value_complex III_ = 0.000790; **Figure 7A**).

**Figure 7.**
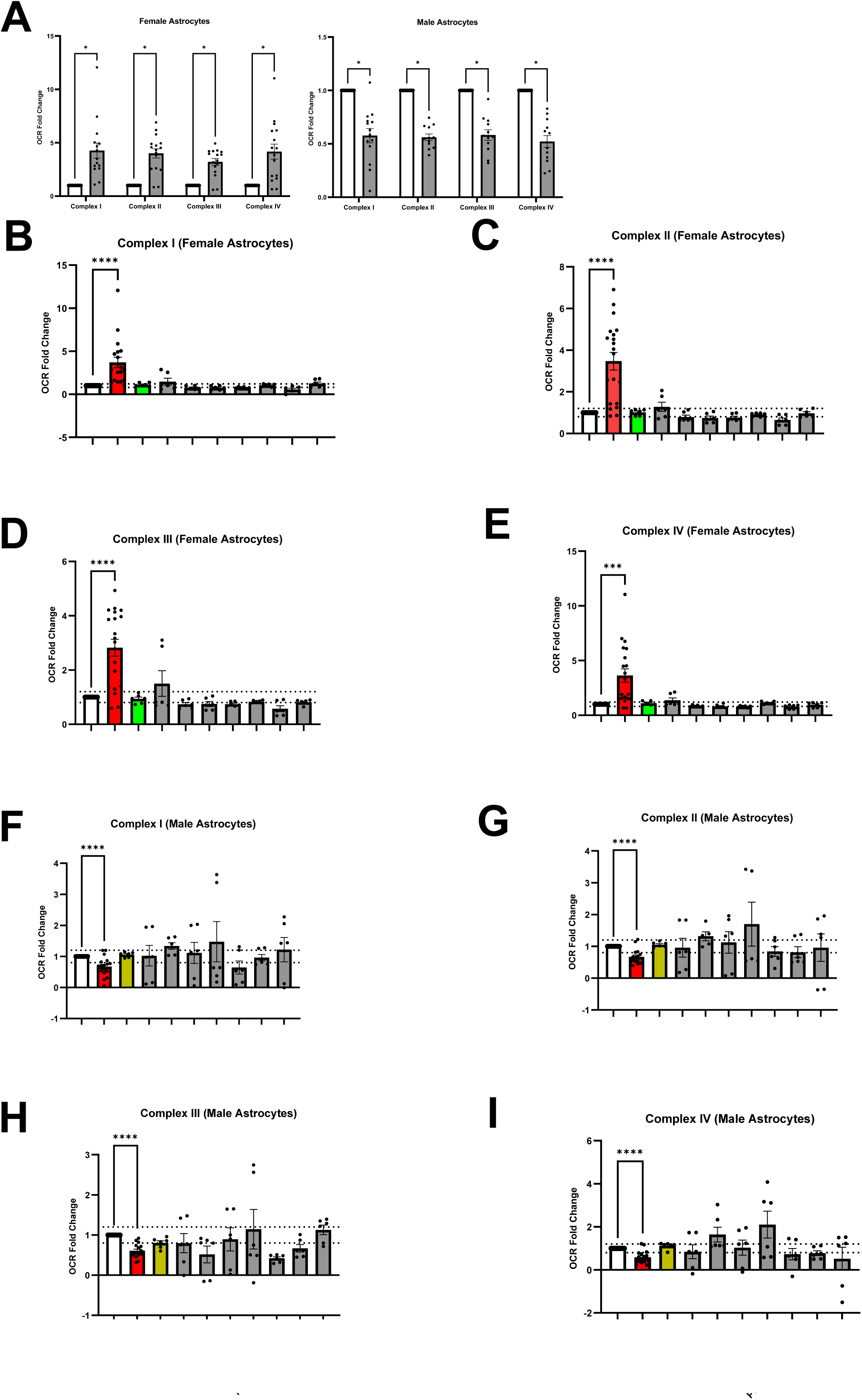

The drug screen in iPSC-derived astrocytes provided distinct, non-overlapping hits from those identified in neurons. In female astrocytes, anthocyanidin, an anthocyanin with reported antioxidant and anti-inflammatory properties was the sole protective hit (**Figure 7B-E**). In male astrocytes a PPARα agonist was the sole protective hit (**Figure 7F-I**). This further underscores the sex- and cell type-specificity of mitochondrial dysfunction in AD.

### Mitochondrial drugs modulate mitochondrial and inflammatory markers in sporadic AD iPSC-derived astrocytes

In female AD iPSC-derived astrocytes, MitoSOX, TMRE, and MitoTracker fluorescence, were not altered (**data not shown**). Our secondary screen confirmed anthocyanidin normalized OCR in female AD astrocytes across multiple independent donors (**Figure 8A**, n=6). Anthocyanidin treatment normalized glial fibrillary acidic protein (GFAP) expression levels in female AD astrocytes (**Figure 8B, C**).

**Figure 8.**
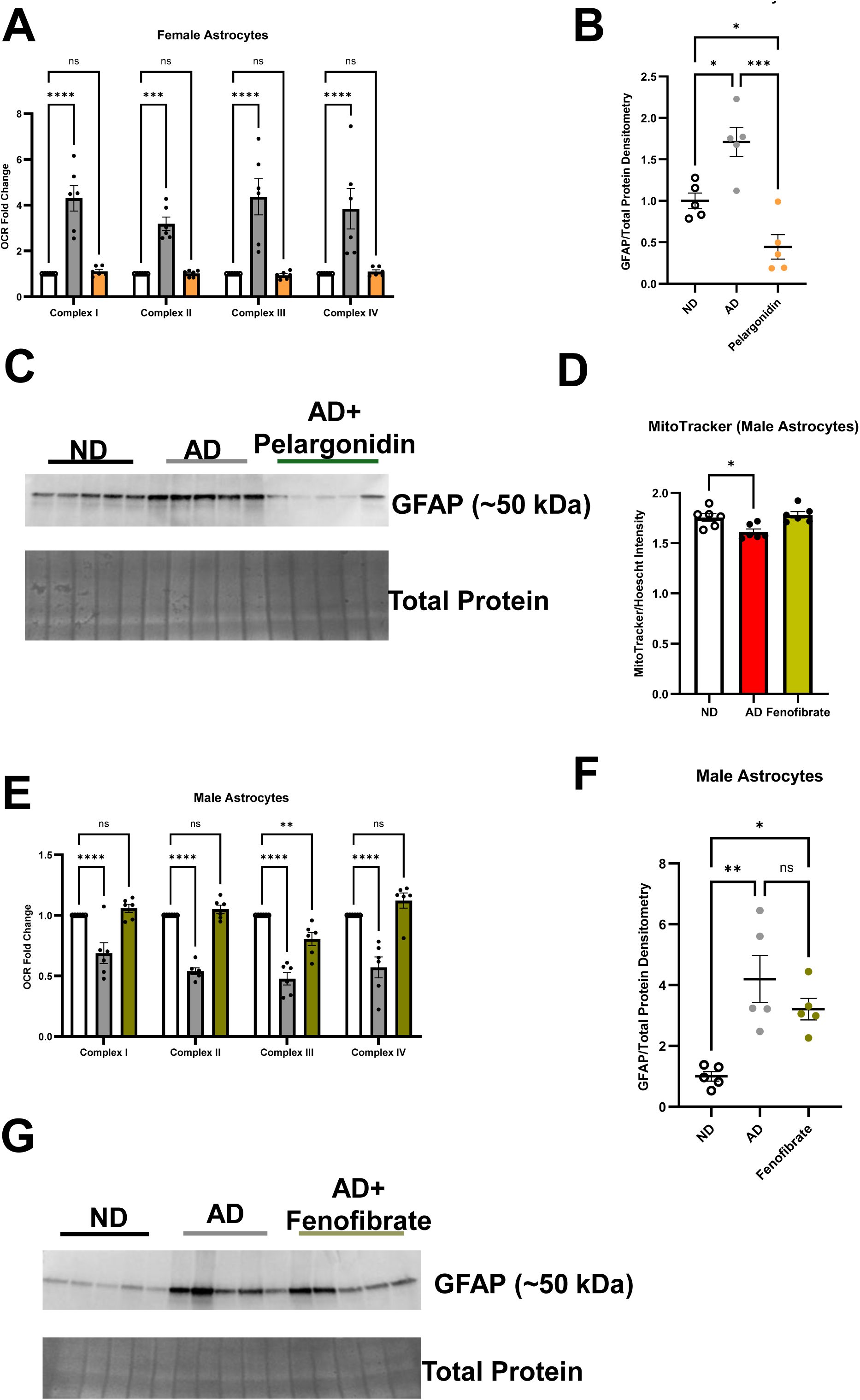

Male AD astrocytes displayed a significant reduction in mitochondrial mass detected by MitoTracker compared to ND controls (**Figure 8D**; Ordinary one-way ANOVA with multiple comparisons to AD, p = 0.0155), suggesting issues with mitochondrial biogenesis or increased mitochondrial turnover in AD astrocytes. This phenotype was rescued by treatment with PPARα agonist (**Figure 8D**), likely due to PPAR agonism. Male AD astrocytes did not show any change in TMRE or MitoSox fluorescence (**data not shown**). Our secondary screen confirmed PPARα agonist normalized OCR in male AD astrocytes across multiple independent donors (**Figure 8E**, n=6). PPARα agonist treatment partially normalized glial fibrillary acidic protein (GFAP) expression levels in male AD astrocytes (**Figure 8F,G**).

### Mitochondrial targeted treatment alters expression of mitochondrial electron transport chain components in iPSC derived astrocytes from sporadic AD **patients.**

Examining mitochondrial ETC subunit abundance identified mostly downregulation across complex I, II, II, IV and V components in female and AD iPSC astrocytes compared to ND neurons (**Figure 9 A-J**). Treatment with anthocyanidin in female AD astrocytes largely normalized expression of ETC components across all complexes (**Figure 9 A, C, E, G, I**). Treatment with PPARα agonist in male AD astrocytes also largely normalized expression of ETC components across all complexes (**Figure 9 B, D, F, H, J**).

### Mitochondrial targeted treatment alters expression of mitochondrial quality control proteins in iPSC derived astrocytes from sporadic AD patients

Anthocyanidin (in female cells) and PPARα agonist (in male cells) normalized mitochondrial dynamics, biogenesis, mitophagy and proteostasis pathway protein expression in female and male AD astrocytes (**Figure 10**). These pathways were either up- or down regulated in AD astrocytes compared to ND astrocytes (**Figure 10**).

### Mitochondrial targeted treatment alters pathway expression in AD astrocytes

AD female astrocytes showed downregulation of translation pathways with upregulation of histone and chromatin modifying pathways (**Figure 11A**). Treatment of female AD astrocytes with anthocyanidin decreased pathways associated with ubiquitination and protein phosphorylation with upregulation of protein sorting and biosynthesis of glycerophospholipid (**Figure 11B**).

Male AD astrocytes showed down regulation of mitochondrial translation and other translation pathways with upregulation of histone and chromatin modifying pathways (**Figure 11C**). PPARα agonist treatment upregulated pathways associated with growth and endocytosis, but downregulated pathways associated with RNA processing and translation when compared to untreated male AD astrocytes (**Figure 11D**). Other pathway changes can be found within **Supplemental Tables 1** and **2**.

### Sex-dependent restoration of mitochondrial respiration by combination treatment

Seahorse analysis revealed distinct sex-dependent alterations in mitochondrial respiratory function in AD co-cultures (**Figure 12A,B**). Female AD co-cultures exhibited elevated OCR relative to ND controls across mitochondrial respiratory complexes (1.44-, 1.94-, 1.85-, and 1.61-fold for Complexes I–IV, respectively, **Figure 12C**). Neither P38 MAPKI nor anthocyanidin alone consistently restored these abnormalities toward the ND state. In contrast, combined P38 MAPKI and anthocyanidin treatment shifted OCR substantially toward the ND state (0.93, 1.23, 1.18, and 1.10 for Complexes I–IV, respectively, **Figure 12C**).

**Figure 12.**
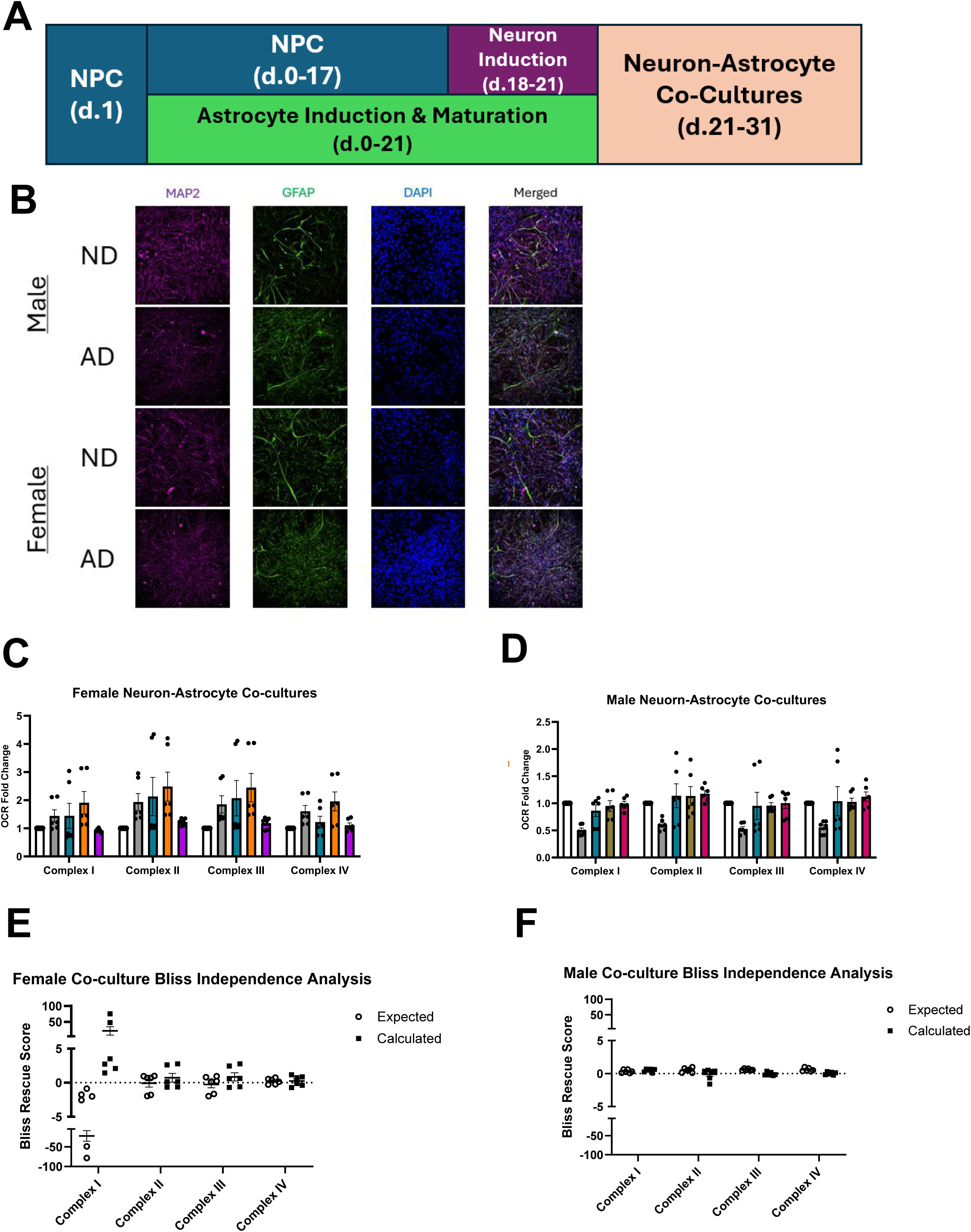

Male AD co-cultures displayed the opposite pattern, with reduced OCR relative to ND controls across all four respiratory complexes (0.51-, 0.61-, 0.53-, and 0.56-fold for Complexes I–IV, respectively, **Figure 12D**). Both P38 MAPKI and PPARα agonist individually shifted OCR toward the ND state. Combined treatment produced further normalization for Complexes I and III, with mean values of 0.99 and 1.00, respectively, while Complexes II and IV showed modest elevations above the ND reference (1.18-and 1.12-fold, respectively, **Figure 12D**). Thus, although the direction of the AD-associated respiratory phenotype differed between sexes, the corresponding combination treatments shifted mitochondrial respiration toward the ND state in both female and male co-cultures.

To evaluate whether the combination effects exceeded those expected from the individual treatments, rescue was normalized according to the distance of each treatment condition from the ND reference and analyzed using a Bliss independence framework. In female co-cultures, the combination produced positive deviations from Bliss independence across all four respiratory complexes, with the largest deviations observed for Complexes I–III (**Figure 12E**). In male co-cultures, combination responses were close to the Bliss-independent expectation for Complexes I and III, whereas modest negative deviations were observed for Complexes II and IV (**Figure 12F**). These findings indicate that the female-directed P38 MAPKI/anthocyanidin combination produced greater than expected restoration of mitochondrial respiratory function, whereas the male directed P38 MAPKI/PPARα agonist combination largely produced responses consistent with independent drug effects.

## Discussion

This study demonstrates that mitochondrial dysfunction in sporadic Alzheimer’s disease (AD) is heterogenous across neural cell types and biological sex. Using patient-derived iPSC neurons and astrocytes, we identified distinct patterns of mitochondrial dysfunction and identified compounds capable of restoring bioenergetic function in a cell-type- and sex-dependent manner. Although both male and female AD neurons displayed impaired respiratory function across mitochondrial complexes I-IV, astrocytes showed more divergent phenotypes with reduced respiration in male AD astrocytes and increased respiration in female AD astrocytes. Consistent with these differences, protective compounds identified from the drug scree were largely cell- and sex-specific. These findings indicate that mitochondrial dysfunction in AD could be characterized as a context-dependent mitochondrial remodeling rather than a uniform reduction in mitochondrial function.

### Cell type and sex shape mitochondrial dysfunction in AD

The shared reduction in complex I–IV respiration observed in male and female AD neurons suggests that impaired oxidative phosphorylation is a robust neuronal phenotype in this cohort. However, the molecular signatures indicate that the cellular response to this dysfunction differs substantially by sex. Male AD neurons showed negative predicted activity of oxidative phosphorylation, respiratory electron transport, complex I biogenesis, complex III assembly, and cristae formation; whereas female AD neurons showed positive predicted activity of these same pathways despite exhibiting reduced respiratory capacity by Seahorse analysis. Thus, pathway activity inferred from protein abundance did not necessarily correspond directly to functional respiratory output.

This apparent disconnect can biologically informative. Increased abundance or predicted activity of mitochondrial pathways in female AD neurons may represent a compensatory response to impaired mitochondrial function rather than increased mitochondrial performance. Consistent with this interpretation, female neurons also showed prominent signatures of mitochondrial protein degradation, RNA degradation, mitochondrial translation, and mitophagy. These findings suggest that female neurons may respond to respiratory dysfunction by increasing mitochondrial turnover and quality-control processes. In contrast, male neurons showed stronger signatures of mitochondrial stress, AMPK/mTOR regulation, autophagy, and oxidative stress responses. Thus, similar functional impairment may arise within different molecular states in male and female neurons.

The astrocyte phenotype further demonstrates why cell type is an important consideration in AD mitochondrial biology. Male AD astrocytes showed reduced respiration across complexes I–IV, accompanied by reduced mitochondrial mass, whereas female AD astrocytes exhibited increased respiration despite proteomic evidence of suppression of oxidative phosphorylation and mitochondrial pathways. The increased respiration in female AD astrocytes therefore should not necessarily be interpreted as enhanced mitochondrial health. It may instead represent a compensatory or stress-associated increase in respiratory activity that occurs despite underlying disruption of mitochondrial maintenance and redox pathways. Together, these findings suggest that astrocytes may respond to AD-associated stress differently from neurons and that sex can further modify this response.

### p38 MAPK inhibition as a potential broad neuronal intervention

Among the neuronal hits, P38 MAPKI emerged as a particularly compelling candidate because its effects extended beyond restoration of OCR. P38 MAPKI improved multiple mitochondrial phenotypes, including respiratory function and oxidative stress, and in male neurons normalized elevated mitochondrial mass. Importantly, treatment was also associated with reductions in secreted Aβ_42_ and pTau-217. The convergence of mitochondrial and canonical AD-associated phenotypes provides evidence that p38 inhibition can influence multiple pathological features within the same cellular system.

The elevated phospho-p38 observed in AD neurons provides additional biological context for this response. P38 MAPKI did not reduce phospho-p38 abundance, which is consistent with its proposed mechanism of inhibiting p38 MAPK substrate phosphorylation rather than preventing upstream MKK3/6-mediated phosphorylation of p38 itself. Thus, persistence of phospho-p38 following treatment does not indicate failure of pathway inhibition. Instead, it is compatible with inhibition occurring downstream of p38 activation.

The reduction in pTau-217 following P38 MAPKI treatment is also consistent with the established ability of p38 signaling to contribute to tau phosphorylation [30]. However, the present findings do not establish that p38 inhibition directly mediates all the observed effects. p38 signaling itself regulates broad networks involved in cellular stress, protein homeostasis, metabolism, and inflammatory responses [31, 32]; and our proteomic analysis demonstrated extensive remodeling of these pathways following P38 MAPKI treatment. Therefore, the effects of P38 MAPKI may reflect coordinated changes across several downstream processes rather than a single linear p38-to-mitochondria mechanism.

Importantly, the proteomic data reinforces this interpretation. P38 MAPKI produced broad remodeling of RNA regulation, protein homeostasis, stress-response, autophagy, and mitochondrial quality-control pathways. These effects were also strongly sex dependent. In female neurons, treatment was associated with increased mitochondrial protein degradation, mitochondrial RNA degradation, and mitochondrial translation relative to AD neurons, whereas male neurons showed stronger activation of broader autophagy-associated pathways. Thus, P38 MAPKI may improve neuronal function within distinct pre-existing molecular states rather than through a single sex-independent mechanism.

### Unique astrocyte responses identify potentially sex-specific therapeutic strategies

The astrocyte screen identified anthocyanidin and PPARα agonist as distinct protective compounds in female and male astrocytes, respectively. The divergent responses are consistent with the broader sex-dependent molecular states observed in these cells.

In male astrocytes, PPARα agonist treatment increased mitochondrial mass and produced pathway-level changes consistent with activation of oxidative-stress defense and PPARα-associated metabolism. Treatment was associated with increased predicted ROS detoxification, NRF2-mediated stress responses, PPARα-dependent lipid metabolism, and PPARα/RXRα signaling relative to untreated AD astrocytes. These changes provide a mechanistic framework consistent with PPARα agonist’s known activity. However, because the mitochondrial respiratory proteome remained substantially different from ND following treatment, the data suggest selective remodeling rather than complete restoration of mitochondrial function.

Anthocyanidin produced a different response in female astrocytes. Its ability to restore OCR occurred without substantial normalization of mitochondrial mass, membrane potential, superoxide production, or the broader mitochondrial proteomic signature. Instead, pathway changes were concentrated in protein regulation, cellular stress, and inflammatory signaling. This raises the possibility that anthocyanidin improves astrocytic respiratory function through regulation of cellular stress or inflammatory pathways rather than direct mitochondrial biogenesis. This interpretation remains provisional, however, because the current data are associative and do not establish a causal link between these pathways and the OCR response.

Instead, the sex-specific astrocyte hits provide an important proof of concept for using cell type and sex as biological variables during drug discovery. Rather than identifying a single compound that produces equivalent effects across all neural cell populations, the screen revealed compounds whose effects corresponded to distinct cellular states.

### Implications for AD drug discovery

These findings have broader implications for mitochondrial-targeted drug development in AD. Mitochondria are often considered a common therapeutic target because mitochondrial dysfunction is observed across multiple cell types and disease stages [18, 33]. However, our findings indicate that the nature of mitochondrial dysfunction can differ substantially according to both cellular identity and sex. A compound that improves neuronal respiration may have little effect on astrocytes, while a compound effective in astrocytes may act through pathways unrelated to those responsible for neuronal rescue.

This heterogeneity may help explain why targeting a broadly defined mitochondrial phenotype can produce inconsistent therapeutic effects. Rather than treating mitochondrial dysfunction as a single disease mechanism, future approaches may benefit from identifying the specific mitochondrial state present within each vulnerable cell population. In this framework, functional phenotyping can be used to identify candidate compounds, while proteomics and complementary molecular assays can help define the cellular context in which those compounds are effective.

The current findings also support a broader view of astrocytes in AD drug discovery. Astrocytes are increasingly recognized as active regulators of neuronal metabolism, redox homeostasis, lipid metabolism, and inflammatory signaling [17, 34, 35]. The distinct astrocyte phenotypes identified here suggest that therapeutic strategies directed toward astrocytic metabolism may complement neuron-focused interventions.

Specifically, the male-specific response to PPARα agonist and female-specific response to anthocyanidin raise the possibility that metabolic and inflammatory pathways could represent different therapeutic entry points across astrocyte populations.

Several limitations should be considered. First, the study was performed using a relatively limited number of donor-derived iPSC lines, and additional donors will be required to establish the generalizability of the observed sex-specific phenotypes. Larger cohorts will also be important for determining whether the identified drug responses represent robust biological effects or donor-specific responses. Second, iPSC-derived neurons and astrocytes do not fully reproduce the aged human brain environment [10]. Reprogramming can reset aspects of cellular aging [36], and the cultures lack many of the multicellular and vascular interactions that influence mitochondrial function *in vivo*.

Third, pathway activation scores derived from proteomic data represent predictions based on protein abundance and curated pathway relationships and should not be interpreted as direct measurements of pathway activity. This is particularly important where proteomic pathway direction and functional OCR measurements are divergent. Targeted experiments will therefore be necessary to determine whether pathways identified by the proteomic analysis are causally responsible for drug-mediated rescue.

Finally, although P38 MAPKI produced the most extensive neuronal rescue phenotype, the current experiments do not establish whether p38 inhibition is sufficient to reproduce all its effects or whether additional targets contribute to the response. Similarly, the mechanisms responsible for the astrocyte-specific effects of anthocyanidin and PPARα agonist remain to be established. Dose-response studies, pharmacological and genetic target-validation experiments, mitochondrial substrate and flux measurements, and validation in independent donor cohorts will be important next steps. Ultimately, testing the leading candidates in appropriate *in vivo* models will be necessary to determine whether the cell- and sex-specific effects observed in iPSC-derived cells translate to the intact brain.

In summary, this study demonstrates that mitochondrial dysfunction in AD is heterogeneous across both cell type and biological sex. Although male and female AD neurons shared a core phenotype of impaired respiratory capacity, their underlying molecular responses differed substantially. Astrocytes showed even greater divergence, with reduced respiration in male AD astrocytes and increased respiration in female AD astrocytes, accompanied by distinct molecular signatures and drug responses.

P38 MAPKI emerged as a lead neuronal candidate because it improved mitochondrial function across multiple readouts and was additionally associated with reduced Aβ42 secretion and pTau-217 abundance. Its effects occurred despite persistent phospho-p38, consistent with inhibition of downstream p38 signaling and were accompanied by broad remodeling of mitochondrial quality-control, stress-response, and proteostatic pathways. In astrocytes, PPARα agonist and anthocyanidin produced sex-specific rescue phenotypes, with PPARα agonist-associated changes consistent with PPARα-linked metabolic and antioxidant signaling and anthocyanidin-associated effects pointing toward cellular stress and inflammatory pathways.

More broadly, the observation that functional rescue frequently occurred without complete normalization of the proteomic phenotype suggests that therapeutic benefit does not necessarily require restoration of the diseased cell to an ND-like molecular state. Instead, effective compounds may improve cellular function by remodeling specific pathological processes within an already altered cellular state.

Together, these findings support a precision approach to mitochondrial drug discovery in AD in which cell type and biological sex are incorporated into target identification and therapeutic evaluation. Rather than if a single mitochondrial phenotype or therapeutic mechanism applies across the brain, future studies should consider the distinct metabolic, stress-response, and proteostatic states of neurons and astrocytes when evaluating candidate interventions.

